# Characterisation of the conformational changes of GlnH that stimulate PknG activity in Mycobacteria and *Corynebacterium glutamicum*

**DOI:** 10.64898/2026.07.02.735984

**Authors:** Hannah L. Tompkins, Silke Roscher, Anna D. Liuzzi, Amanda K. Chaplin, Russell Wallis, Helen M. O’Hare

## Abstract

GlnH is an amino acid binding protein that senses aspartate to regulate metabolism via the PknG pathway in diverse Actinobacteria. Information about ligand occupancy of periplasmic GlnH is conveyed to PknG via an uncharacterised transmembrane protein GlnX. This pathway is important in the virulence of *Mycobacterium tuberculosis*, and in regulating valuable industrial fermentations by *Corynebacterium glutamicum*. GlnH has a “Venus flytrap”-like structure, comprising two lobes that surround the ligand aspartate. However, the conformational changes that allow GlnH to initiate this signalling pathway are unknown. To address this question, we produced GlnH from pathogens *M. tuberculosis* and *Mycobacterium marinum* and non-pathogens *Mycobacterium smegmatis* and *C. glutamicum* and used X-ray crystallography and cryo-EM to determine their structures. The results show that amino acid specificity is conserved in all homologues. However, GlnH from Mycobacteria was monomeric and bound aspartate with nanomolar affinity, whereas GlnH from *C. glutamicum* bound aspartate with micromolar affinity and dimerised upon binding. Whilst GlnH of the non-pathogens was stable at neutral pH, GlnH from the pathogens was most stable at acidic pH, reflecting the environment of host phagosomes. Structures were determined for all homologues, but only *M. smegmatis* GlnH crystallised in both unbound (Apo) and Asp-bound forms. GlnH has an open structure with a cleft between the lobes to permit access to aspartate. The Asp-bound structure is more compact with the lobes locked together, completely enclosing the ligand. AlphaFold was used to design mutations to disrupt the predicted GlnH-GlnX interface, and these variants failed to complement the metabolic defect of *glnX* knockout in *M. smegmatis*, supporting the predicted complex and suggesting how the GlnH conformational change is transmitted GlnX to initiate signalling.

**Author summary:** Bacteria sense their environment and respond to changes using a variety of sensors and regulators. We investigated a widely conserved sensor that allows *Mycobacterium tuberculosis* to detect amino acids in order to regulate its metabolism in different environments within the human body. A key requirement of any sensor is the ability to change shape in reponse to its stimulus. We have determined the structures of the sensor in the presence and absence of amino acid to identify the changes in shape and how these could be passed into the bacterial cell to change its behaviour. We used four related organisms to understand how sensing differs between pathogens and non-pathogens.

## Introduction

The Protein kinase G (PknG) pathway consists of a 3-gene operon (*glnX*, *glnH*, *pknG*) and the kinase substrate and effector protein GarA (1). This pathway is widely conserved in the *Corynebacteriales* where it controls glutamate entry into the TCA cycle, and is important for virulence of *Mycobacterium tuberculosis* (2), and for amino acid over-production by industrial *Corynebacterium glutamicum* (3).

In the presence of amino acids, PknG phosphorylates GarA, which promotes glutamate catabolism (2). GlnX and GlnH link amino acid availability to kinase activity: GlnH is a lipoprotein sensor that is exposed to external nutrients and can bind amino acids, while GlnX is a transmembrane protein that is proposed to detect occupancy of GlnH and transduce this information across the membrane to stimulate cytoplasmic PknG (1).

Apart from its involvement in metabolic regulation, some studies have suggested that PknG of *M. tuberculosis* (and potentially other pathogenic Actinobacteria) may have a direct role in modulating macrophage function, potentially involving secretion of PknG (4–6). It is not known whether GlnH and GlnX would be required to recruit PknG to the membrane and/or promote its autophosphorylation for secretion. It is possible that PknG (and potentially GlnH and GlnX) may have diverged during the evolutionary specialisation of pathogenic mycobacteria.

GlnH is a member of the large and diverse family of substrate binding proteins (SBP) found in both prokaryotes and eukaryotes. These proteins have two main roles: ligand-sensing, for example to activate histidine kinases in bacteria or as part of human glutamate receptors, or facilitating ligand uptake through ABC transporters (7). Crystal structures of *M. tuberculosis* GlnH in complex with amino acids have explained its specificity for aspartate, glutamate and asparagine (1). GlnH consists of a large and small lobe, joined by two beta strands, with the amino acid ligand bound in between the lobes. Protein conformational change is a requirement for a sensor but the nature of this changes has not been addressed because GlnH only crystallised in the presence of amino acids (1), and is difficult to predict since SBPs are heterogeneous in their response to ligand: some “open” by less than 2 Å in the absence of ligand, while others hinge open by more than 20 Å ({Berntsson, 2010 #37}).

To address the molecular mechanism of ligand sensing by GlnH, and potentially shed light on any divergence of function, we produced recombinant GlnH from two pathogens: *M. tuberculosis* and *Mycobacterium marinum*, and two non-pathogens: *Mycobacterium smegmatis* and *C. glutamicum*. Compared to *M. tuberculosis* (GlnH_Mt_), the homologues share 89% (*M. marinum*, GlnH_Mm_), 74% (*M. smegmatis*, GlnH_Ms_) and 43% (*C. glutamicum*, GlnH_Cg_) amino acid identity (supplementary figure 1). These proteins were compared for ligand-specificity and affinity, and investigated for the influence of aspartate on protein stability and structure.

## Results

### Production of recombinant GlnH and characterisation of ligand binding

GlnH from *M. tuberculosis* (GlnH_Mt_) and *C. glutamicum* (GlnH_Cg_) were previously found to bind a variety of amino acids, with highest affinity for aspartate: K_D_ of 230 nM and 264 μM respectively (8, 9). Using a similar strategy, the homologues from *M. marinum* (GlnH_Mm_) and *M. smegmatis* (GlnH_Ms_) were produced in soluble form by replacing the N-terminal lipobox motif with a His_6_ tag. Aspartate was identified as the optimal ligand for both GlnH_Mm_ and GlnH_Ms_ as it caused the greatest increase in melting temperature (figure 1 A&B). Other amino acids caused smaller increases (glutamate, asparagine, cysteine and histidine) or no significant change (glutamine and glycine). The affinity for aspartate was determined by isothermal calorimetry (figure 1 C&D), with both GlnH_Mm_ and GlnH_Ms_ having K_D_s in the nanomolar range (table 1), similar to GlnH_Mt_ but >100-fold lower than that of GlnH_Cg_, suggesting that sub-micromolar K_D_ for aspartate may be a general feature of mycobacterial GlnH.

**Figure 1.**
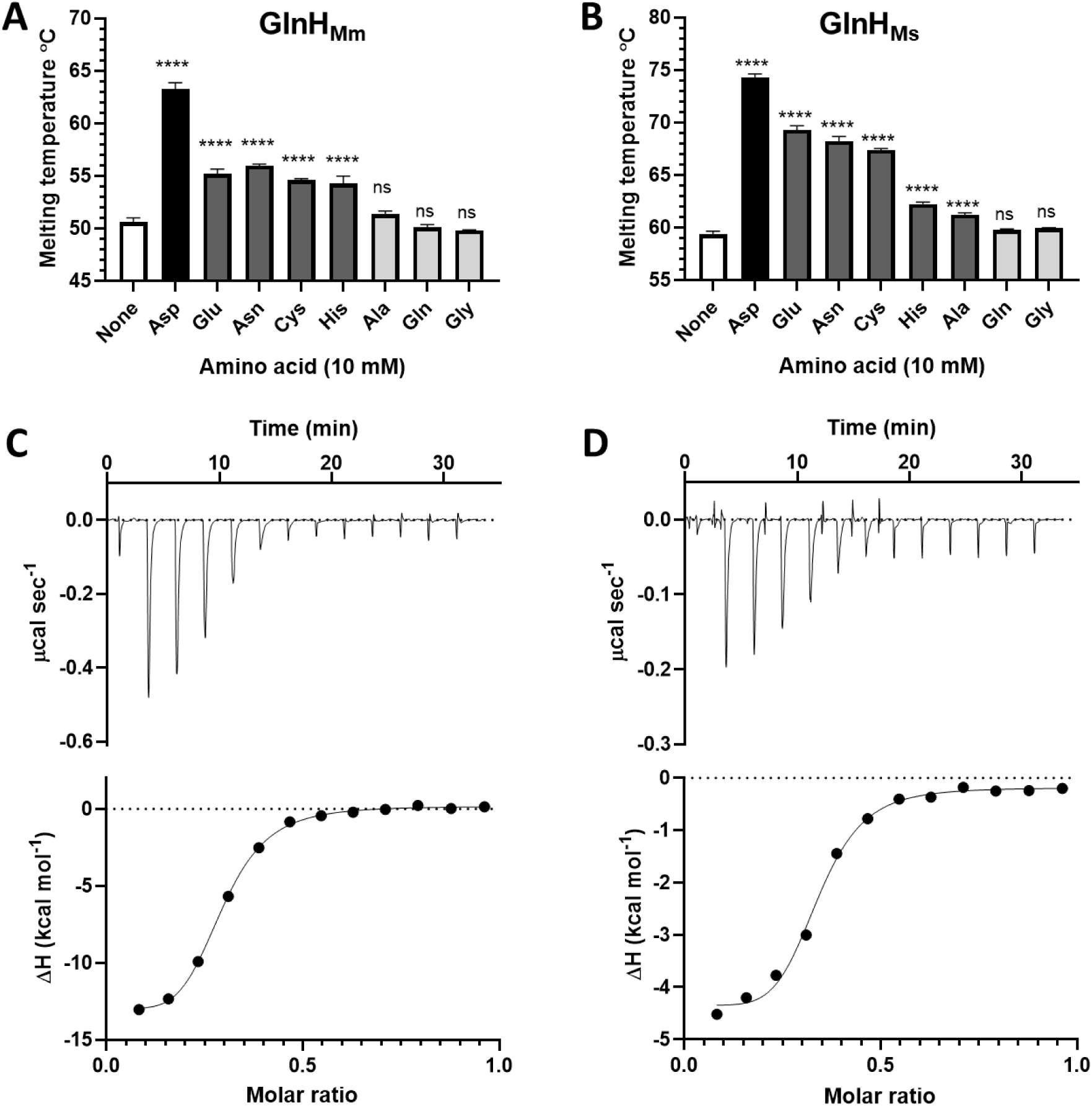
GlnH from pathogens and non-pathogens bound Asp with high affinity (A&B) Binding of amino acids to GlnH_Mm_ (A) and GlnH_Ms_ (B) was measured by thermal melt shift assay. Asp caused the greatest increase in thermal stability and Glu, Asn, Cys and His also caused a significant increase in melting temperature (****, p<0.0001), whereas Gln and Gly did not (ns, p>0.05). Ala stabilised GlnH_Ms_ but not GlnH_Mm_. (C&D) Binding of Asp to GlnH_Ms_ (C) and GlnH_Mm_ (D) measured by isothermal calorimetry (ITC). Representative traces are shown.

**Table 1.**
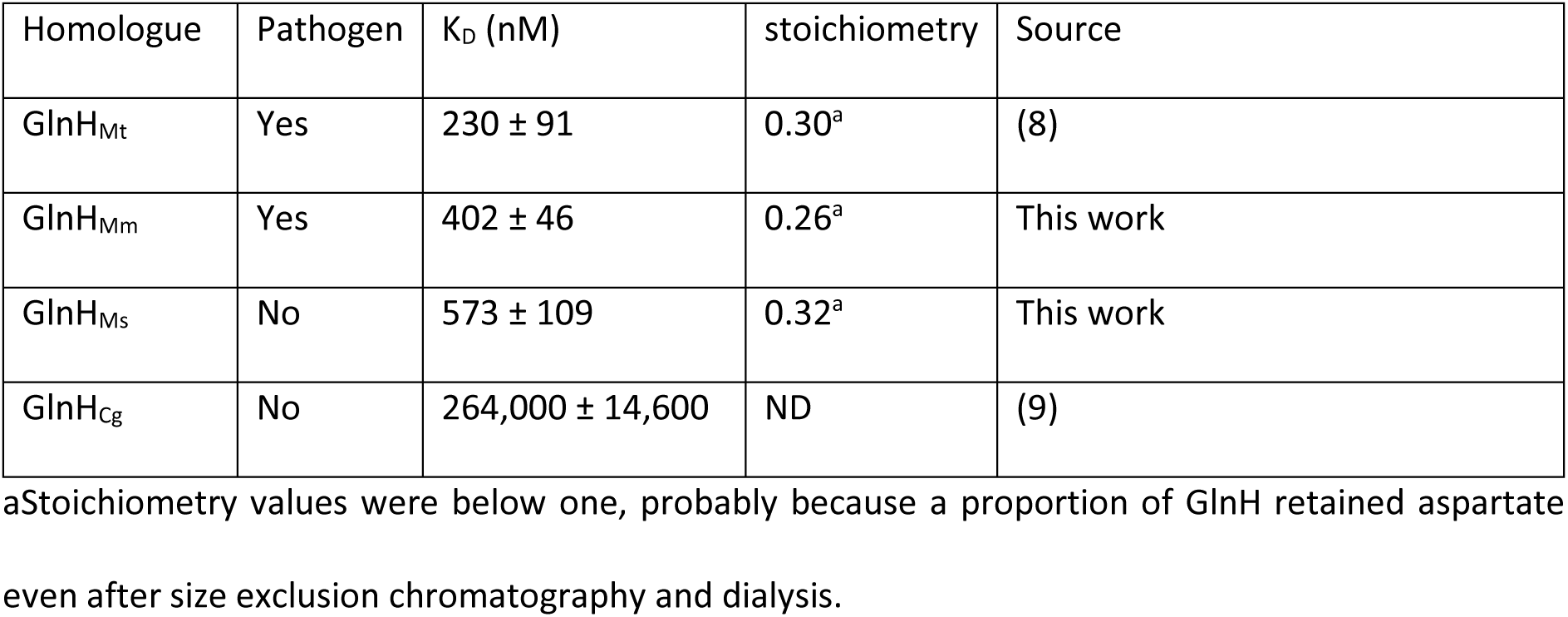

| Homologue | Pathogen | K<sub>D</sub> (nM) | stoichiometry | Source |
| --- | --- | --- | --- | --- |
| GlnH<sub>Mt</sub> | Yes | 230 ± 91 | 0.30<sup>a</sup> | (8) |
| GlnH<sub>Mm</sub> | Yes | 402 ± 46 | 0.26<sup>a</sup> | This work |
| GlnH<sub>Ms</sub> | No | 573 ± 109 | 0.32<sup>a</sup> | This work |
| GlnH<sub>Cg</sub> | No | 264,000 ± 14,600 | ND | (9) |

<sup>a</sup>Stoichiometry values were below one, probably because a proportion of GlnH retained aspartate even after size exclusion chromatography and dialysis.

### pH-dependence of thermal stability of GlnH homologues

Previous screening of crystallisation conditions revealed that GlnH_Mt_ was most soluble at lower pH (1). The pH of the mycobacterial periplasm likely matches that that of external environment, due to transporters in the outer mycomembrane (10, 11). The pH range in natural niches during the infection cycle of pathogens ranges from 7.6 (cytoplasmic) to 4.5 (maturing phagosome) (12). The thermal melt assay confirmed that GlnH_Mt_ was not only folded at low pH (4.5), but it was significantly more stable at low than neutral pH (figure 2A). This acid-stabilisation was only apparent in the presence of aspartate, and was shared by GlnH_Mm_, albeit to a lesser extent (figure 2B). By contrast, GlnH_Ms_ and GlnH_Cg_ from non-pathogens were less stable at low pH than at neutral pH, even in the presence of aspartate.

**Figure 2.**
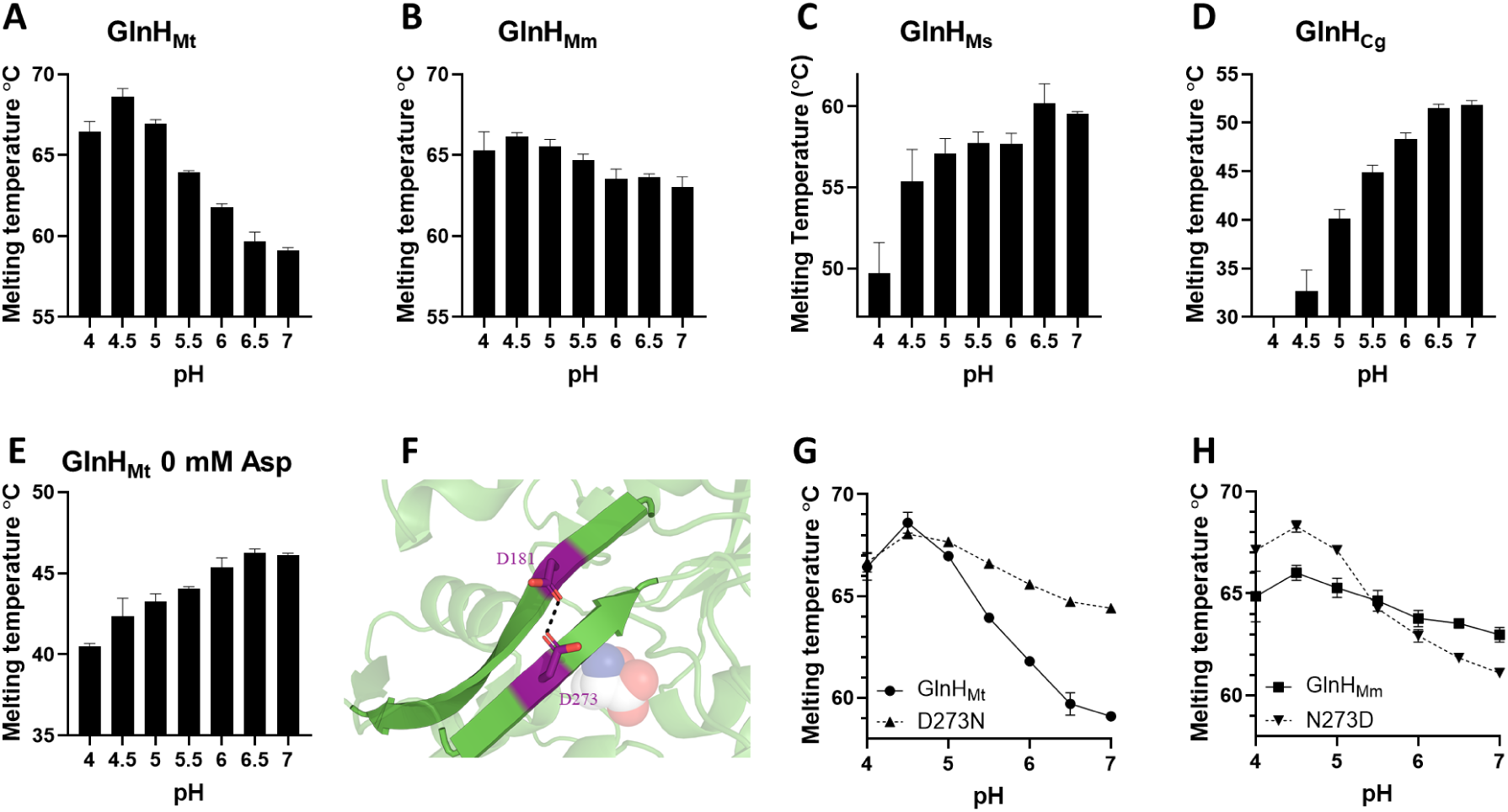
GlnH from pathogens was more stable at acidic pH. (A-D) GlnH from pathogens GlnH_Mt_ (A) and GlnH_Mm_ (B) had a significantly higher melting temperature at pH 4.5 than pH 7, whereas GlnH from non-pathogens GlnH_Ms_ (C) and GlnH_Cg_ (D) had a lower melting temperature at pH 4.5 than pH 7 (p<0.0001). (E) Acid-stabilisation was not observed for unliganded GlnH_Mt_. (F) GlnH_Mt_ has an unusual Asp-Asp hydrogen bond (D181 and D273), whereas the equivalent residues in GlnH_Mm_ are Asn-Asp. The ligand aspartate is shown as a space-filling model. (G) Engineering Asn-Asp into GlnH_Mt_ by the mutation D181N enhanced its stability at neutral pH. (H) Engineering Asp-Asp into GlnH_Mm_ by the mutation N273D enhanced its stability at low pH and reduced its stability at neutral pH.

The crystal structure of GlnH_Mt_ showed an unusual Asp-Asp hydrogen bond in the β-sheet connecting the two lobes (figure 2F), that relies on protonation of one of the Asp side chains at low pH. The equivalent residues in GlnH_Mm_ are Asn-Asp, that could form a hydrogen bond at neutral pH (supplementary figure 1). Site directed mutagenesis was used to investigate the role of this hydrogen bond: introducing Asn-Asp into GlnH_Mt_ increased its stability at neutral pH to resemble GlnH_Mm_ (figure 2G), whereas introducing Asp-Asp into GlnH_Mm_ increased its stability at low pH and decreased its stability at neutral pH to resemble GlnH_Mt_ (figure 2H).

Conformational change is a requirement for a sensor protein, and is also required for ligand binding and release from GlnH since the ligand is solvent inaccessible in the crystal structure (supplementary figure 2). Limited trypsin proteolysis was used to measure differences between Asp-bound and Apo-GlnH. In the absence of aspartate, Mycobacterial GlnH was significantly more susceptible to proteolysis (figure 3), indicative of alternative conformation(s) that are more open or flexible, exposing trypsin cleavage sites. Titration of Asp with a fixed concentration of trypsin confirmed the high affinity of Mycobacterial GlnH for Asp. GlnH_Cg_ was fully digested at all tested concentrations of Asp.

**Figure 3.**
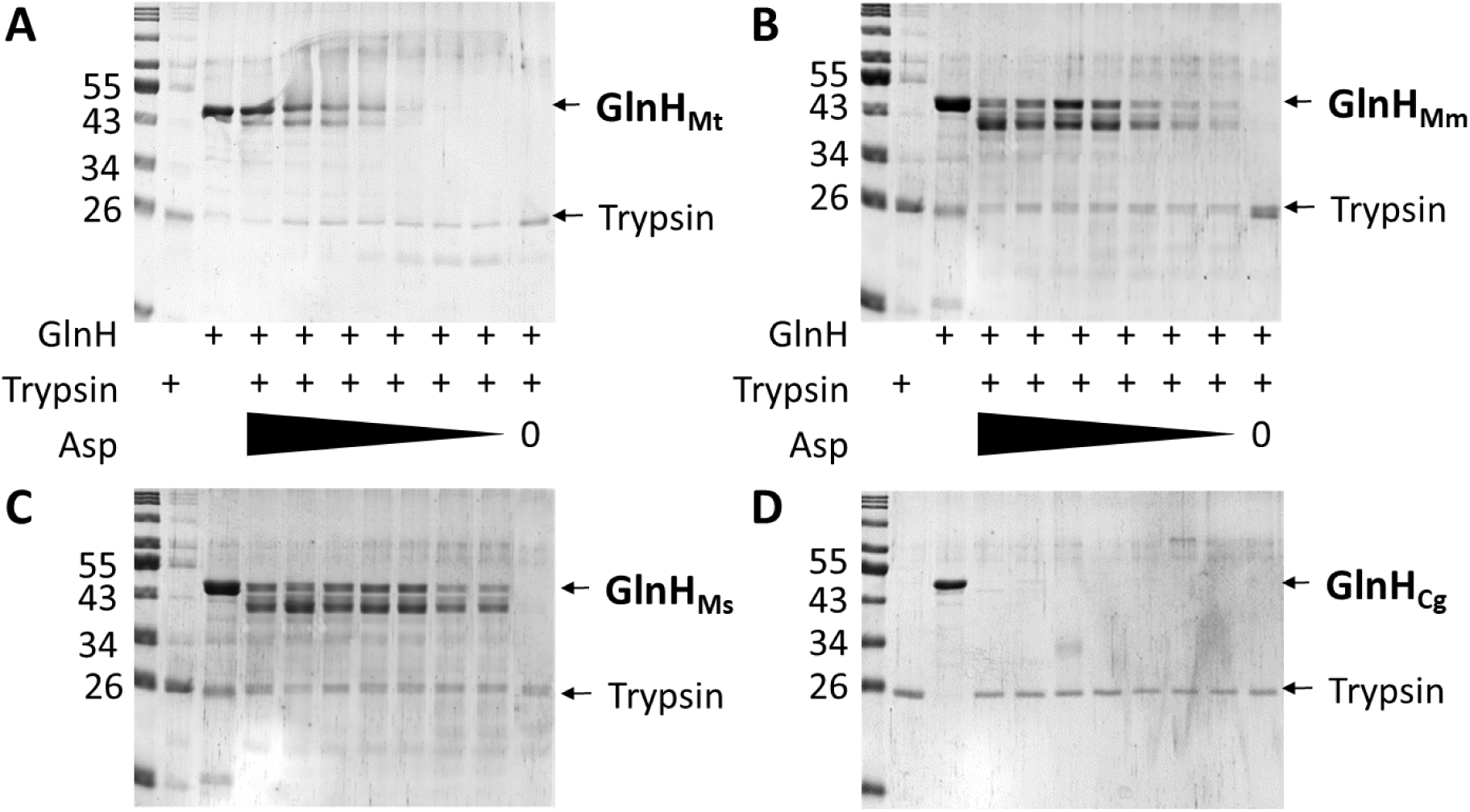
Asp-bound GlnH was more resistant to proteolysis. GlnH_Mt_ (A), GlnH_Mm_ (B), GlnH_Ms_ (C) and GlnH_Cg_ (D) were digested with trypsin in the presence of decreasing concentrations of Asp (5-fold dilutions from 10 mM – 0.64 μM). Protein size markers are in kDa. Bands corresponding to full length GlnH and trypsin are marked with arrows. Gels are representative of multiple independent experiments.

### Aspartate-dependent dimerisation of GlnH_Cg_

All four GlnH proteins were purified as monomers, but GlnH_Cg_ uniquely formed homodimers upon addition of Asp, as observed by size exclusion chromatography and mass photometry (figure 4). After failed attempts at crystallisation, the structure of dimeric GlnH_Cg_ was determined by single particle cryo-EM with an overall resolution of 3.4 Å (figure 5 and supplementary figures S3-S5). Homodimers were parallel and symmetrical (C2 symmetry). Dimerisation involved the C-terminus, both lobes and the interconnecting β-sheet, with the C-terminus domain-swapping and with the ligand binding pocket outermost (figure 5A&B). The buried area of 2007 Å^2^ per monomer was predominantly hydrophobic (figure 5C). To understand whether homodimerisation might be a general feature of GlnH, we examined whether the residues involved in the interface are conserved within the Corynebacterium and Mycobacterium (Supplementary figure S6). The C-terminal residues primarily responsible for the interface are not conserved in *Mycobacterium*, nor in other *Corynebacterium* except for the closest homologue examined, explaining the absence of dimerisation of Mycobacterial GlnH. The overall fold of GlnH_Cg_ resembles the fold of GlnH_Mt_, with the ligand sandwiched between the two lobes (Figure 5B). Any movement of the lobes to release ligand would create a clash and disrupt the dimer interface, explaining why dimerisation is ligand-dependent. Notably the positions of putative Asp-binding residues were similar to their counterparts in the crystal structures of GlnH_Ms_, Gln_Mm_ and GlnH_Mt_ with relatively good density surrounding the Asp binding site, including the Asp ligand itself (Figure 5B). A lower resolution map was obtained from a grid of monomeric GlnH_Cg_ prepared in the absence of aspartate (4.6 Å), and refinement showed that the overall fold of the monomer is similar to that of the homodimer (figure 5D). However, the large lobe is rotated by ∼10 degrees with respect to the small lobe along the long axis, leading to displacement of the secondary structure elements by 3-3.5 Å (figure 5E). The resulting structure is incompatible with dimerisation. The resolution was insufficient to determine whether these changes permit ligand access.

**Figure 4.**
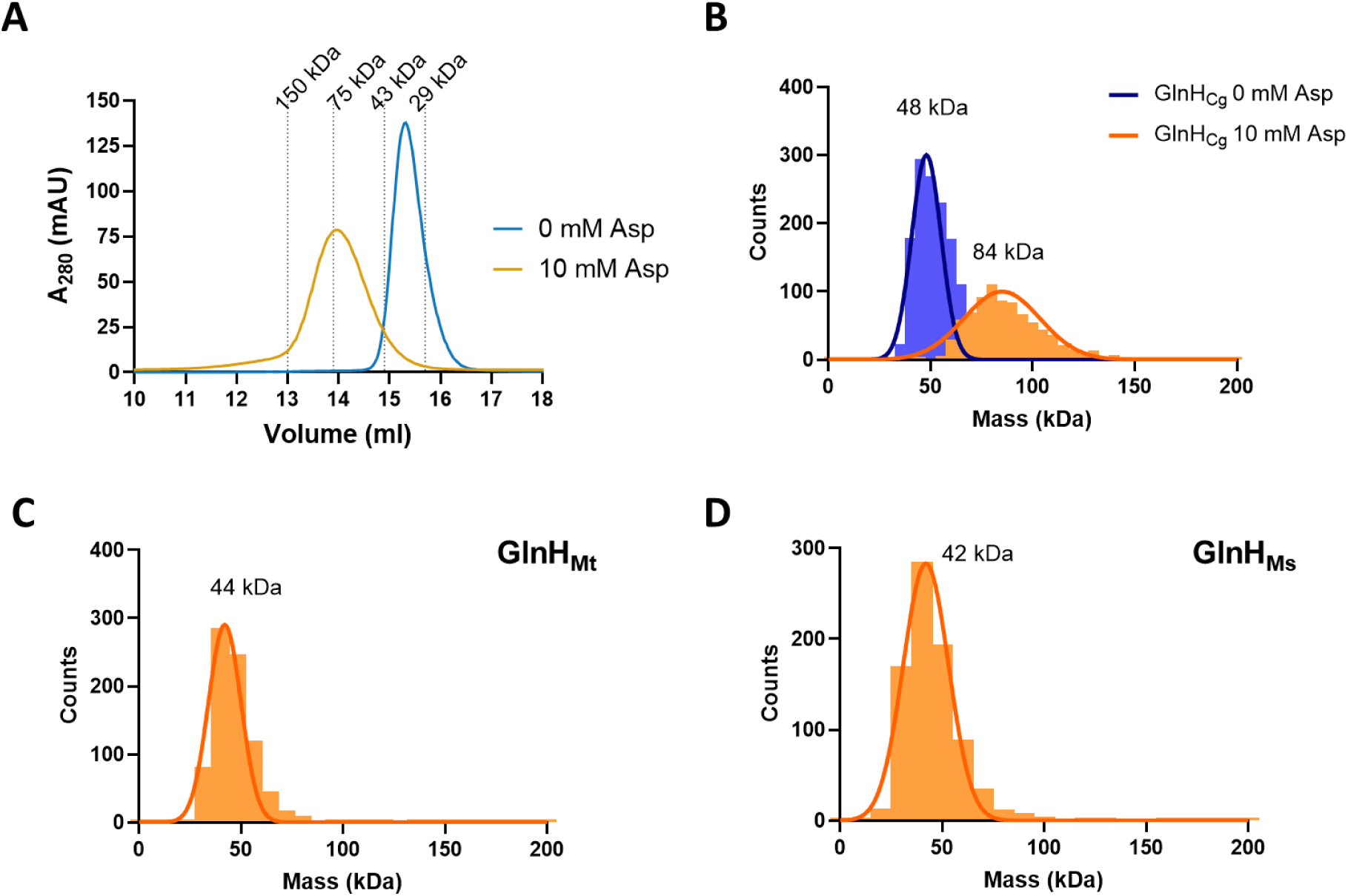
GlnH_Cg_ dimerised in the presence of Asp, whereas Mycobacterial GlnH was monomeric. (A) Size exclusion chromatography of GlnH_Cg_ showed mobility corresponding to monomers at 0 mM Asp and dimers at 10 mM Asp. Dashed lines indicate the mobility of protein size standards. The expected molecular mass of monomers was 37.0 kDa. (B) Mass photometry of GlnH_Cg_ revealed a peak of 48 kDa corresponding to monomers (sigma 7.2 kDa, 1125 counts, skewness 0.000), or 84 kDa corresponding to dimers when Asp was included (sigma 16. 4 kDa, 794 counts, skewness 0.000). No larger oligomers were observed. (C&D) Mass photometry of GlnH_Mt_ (C) and GlnH_Ms_ (D) measured peaks of 44 kDa and 42 kDa respectively corresponding to monomers in the presence of Asp (Sigma 8.1 kDa, 771 counts (93%), skewness 0.000 and sigma 11.1 kDa, 776 counts (94%), skewness 0.000). The expected molecular masses were 35.4 kDa and 34.0 kDa.

**Figure 5.**
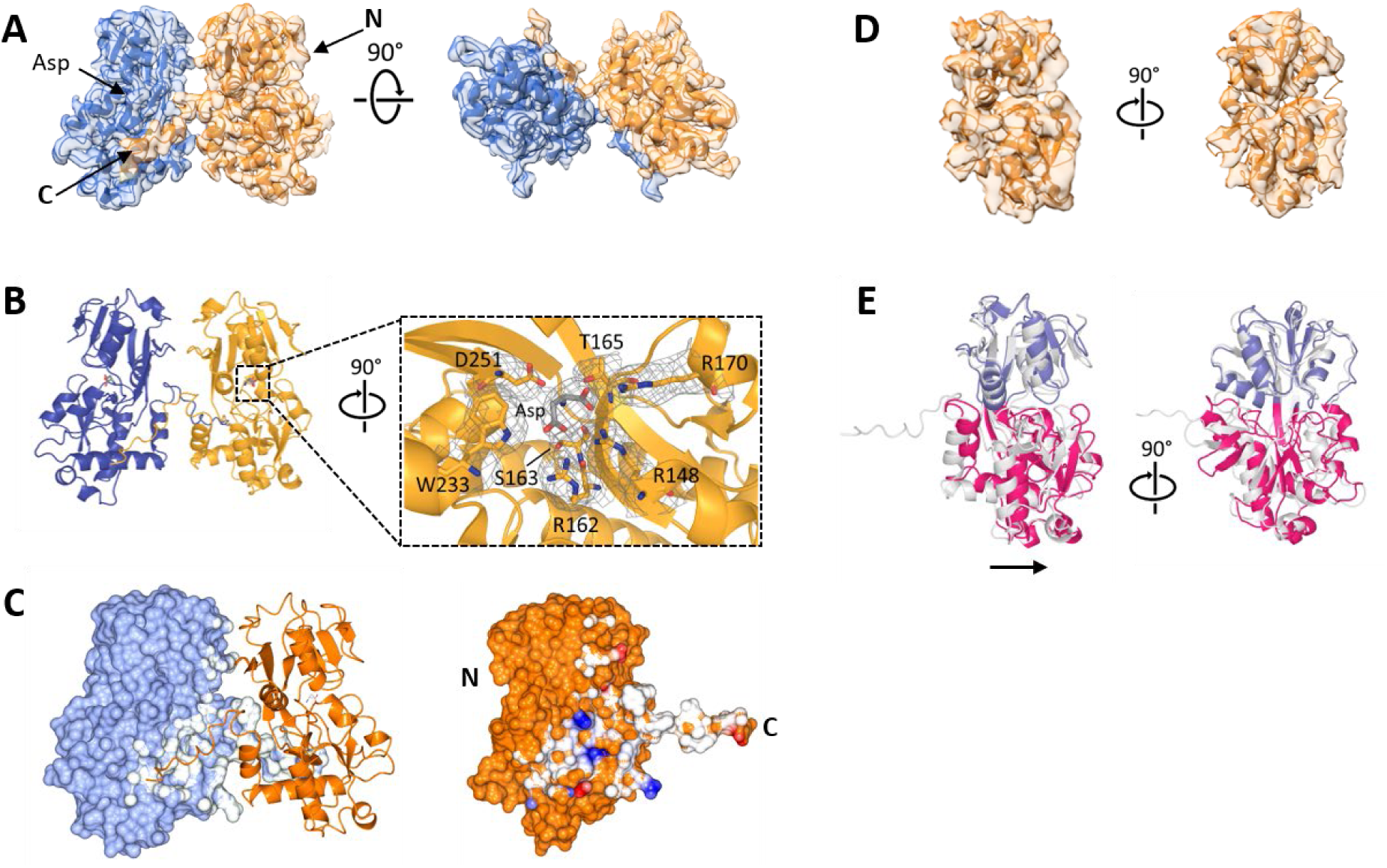
Structures of monomeric Apo-GlnH and dimeric Asp-bound GlnH of *C. glutamicum* by single particle cryo-EM. (A) Cryo-EM map and model of aspartate-bound GlnH_Cg_. Homodimers are symmetrical, with the C-terminus of each protomer swapping over to bind to its partner. (B) The ligand is sandwiched between the two lobes. The β-sheet linking the lobes is towards the dimer interface and the ligand binding site faces outward. Inset: putative ligand-interacting residues are shown with the map at 2 σ. (C) The dimer interface (white) involves both lobes, the connecting β-sheet and the C-terminus, and is predominantly uncharged. The right hand view is rotated to show the whole interface and coloured by electrostatic potential. (D) Cryo-EM map and model of monomeric Apo-GlnH_Cg_. (E) Alignment of the smaller lobe (blue) of Apo (pink and blue) vs aspartate-bound (white) GlnH_Cg_ reveals that removal of aspartate causes rotation of the lobes relative to one another about the long axis. The arrow indicates the direction of displacement.

### Aspartate-induced conformational changes of Mycobacterial GlnH

Aspartate-bound crystal structures were obtained for GlnH_Ms_ and Gln_Mm_, even when aspartate was omitted from purification and crystallisation buffers, likely reflecting the high affinity for and prevalence of aspartate. Structures were similar to that of GlnH_Mt_ reported previously (figure 6). Figure 6B shows the binding site of GlnH_Ms_. The amine and carboxyl groups of the aspartate ligand are stabilized by polar contacts with the side chains of Thr158, Arg 163 and Asp244 and the side-chain carboxyl group of the aspartate interacts with the side chains of Thr156, Arg141 and Trp226. All six residues are conserved in the four homologues described here, with Thr 156 replaced by a serine residue in GlnH_Cg_ and Thr158 replaced by a serine in GlnH_Mt_.

**Figure 6.**
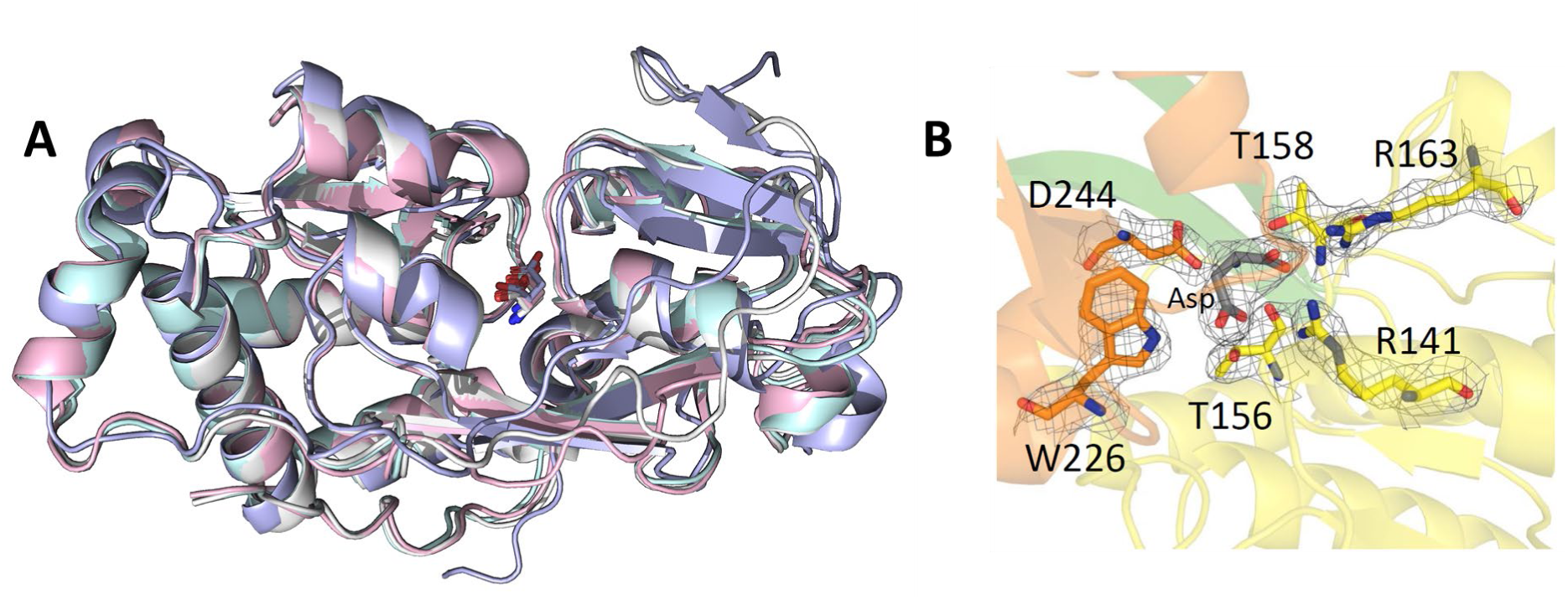
Asp-bound structures of GlnH were similar between species. (A) Crystal structures of GlnH_Ms_ (pink) and GlnH_Mm_ (cyan) were determined and aligned with the cryo-EM structure of dimeric GlnH_Cg_ (purple) and the published crystal structure of GlnH_Mt_ (white; 6H1U). The overall structure and position of the ligand were similar (R.M.S.D. values of 0.34, 0.54 and 0.93 Å from 6H1U respectively). Structures of each species were aligned, omitting eight residues from the C-terminus of GlnH_Cg_ (involved in dimerisation). (B) The ligand binding site of GlnH_Ms_. Residues forming polar contacts with the ligand are shown as sticks. The electron density map is displayed on the Asp ligand and the Asp-binding residues (2Fo − Fc map contoured at 1.6 σ). The lobes are shown in orange (small), yellow (large), with the connecting beta sheet in green.

Interestingly, one of four copies of GlnH_Mm_ in the asymmetric unit was substantially different from the other three, likely reflecting changes induced by crystal contacts. Although, it still bound to Asp, the two lobes were further apart, providing a first clue to the changes that are likely to occur upon release of the ligand (figure 7). All polar contacts were retained with the Asp ligand with the exception of Arg147, which folded back away from the ligand to make a hydrogen bond with the main chain. Moreover, the ligand became accessible to solvent via a narrow channel.

**Figure 7.**
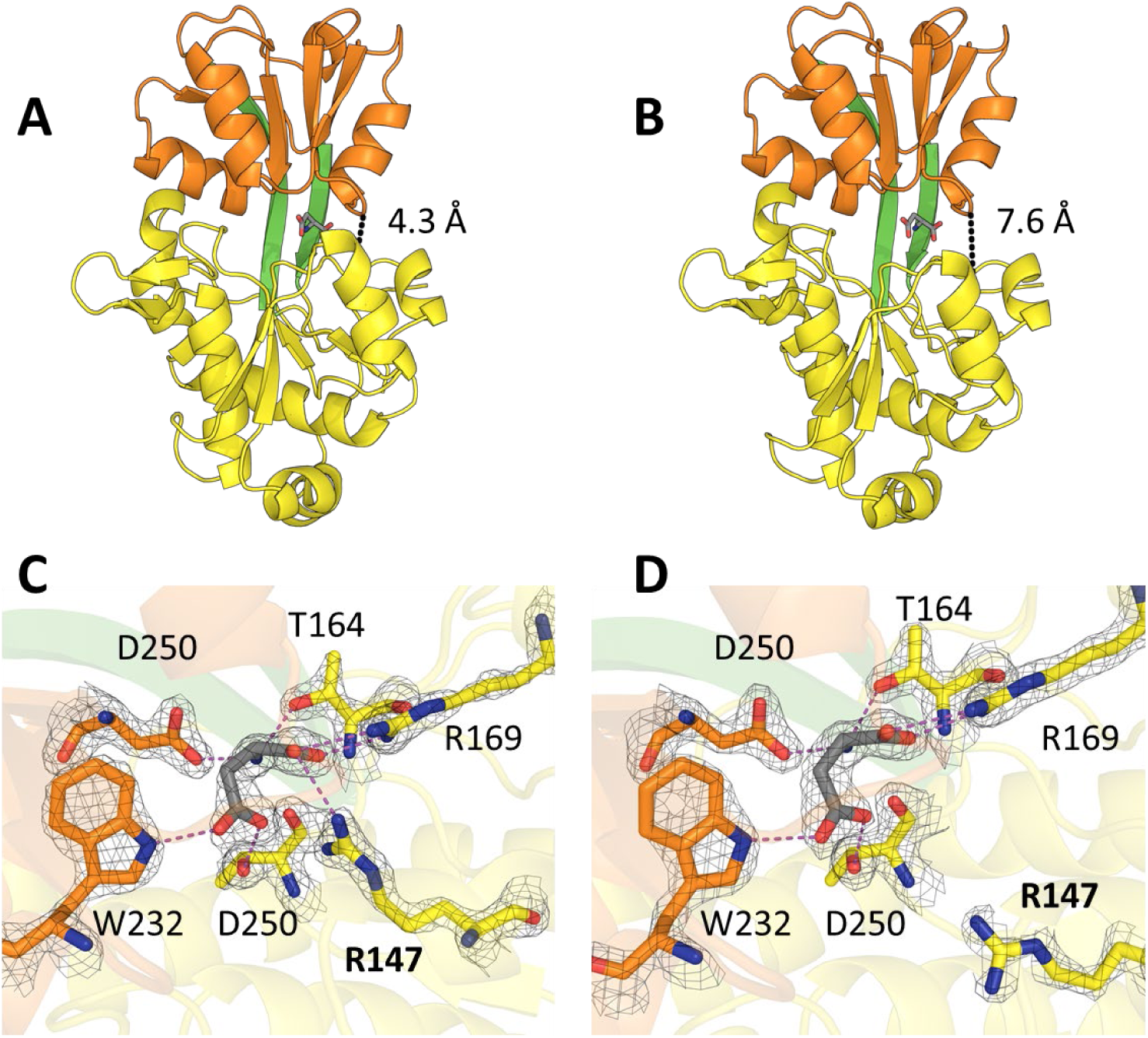
Closed and partially open structures of GlnH_Mm_. (A) Three of the four molecules in the asymmetric unit of GlnH_Mm_ had closed structures in which the small and large lobes were in close contact. The small lobe is orange and the large lobe is yellow. The ligand aspartate is grey and the distance between the lobes is 4.3 Å (dashed line). (B) The fourth molecule of GlnHMm had a greater distance between the lobes (7.6 Å). (C & D) The residues contacting the ligand aspartate were the same in both conformations, except for R147, which contacted the ligand in the closed structure (C) but not in the partially open structure (D).

Given the propensity of GlnH to crystallise bound to aspartate, even after extensive dialysis, we attempted an alternative strategy to characterise the Apo-form by reducing the affinity of GlnH for its ligand. Two site directed mutants of GlnH_Ms_ were designed: a double mutant Thr156Ala Thr158Ala (GlnH_TA_), which lacks two hydrogen bonds to the ligand, and a single mutant Asp244Phe (GlnH_DF_) in which the phenylalanine blocks the ligand binding site. Both variants were folded, as seen by cooperative thermal denaturation, with a similar melting temperature to the wild type protein, but had reduced binding to aspartate and glutamate (figure 8). These variants crystallised readily and diffracted to high resolution (1.64 and 1.60 Å), to reveal the “Apo” conformation of GlnH, with solvent access to the Asp-binding site via a pronounced cleft between the two lobes. To obtain the Apo structure of the wild type, crystals of GlnH_TA_ were used for streak seeding of GlnH_Ms_ and the resulting crystals were used to determine the structure at 1.58 Å resolution (fig 9).

**Figure 8.**
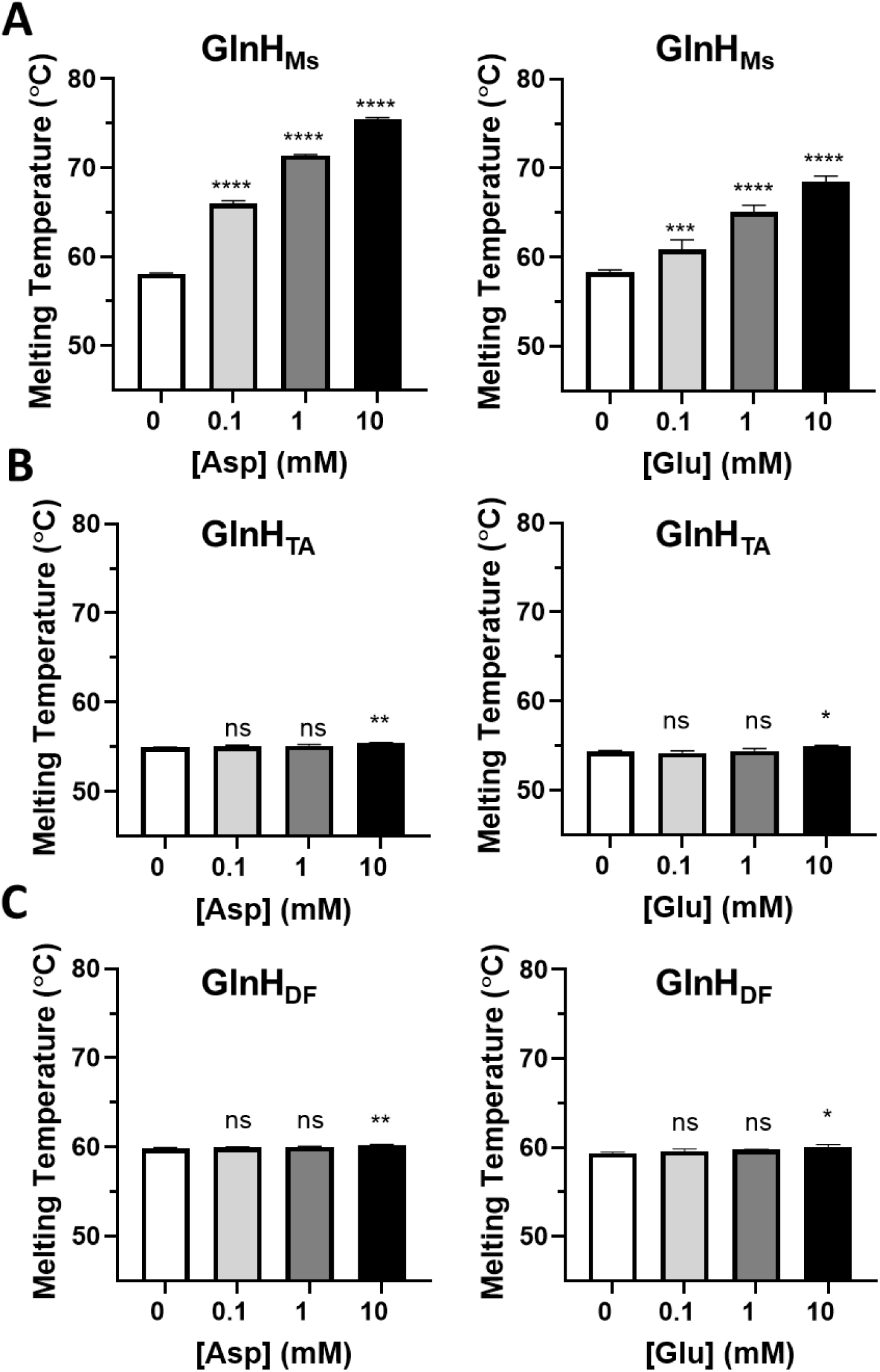
Site directed mutants were designed to reduce binding of Asp to GlnH_Ms_. (A) Wild type GlnH_Ms_ showed increasing melting temperature with increasing concentrations of aspartate or glutamate. (B & C) GlnH_TA_ (T156A T158A) and GlnH_DF_ (D244F) only showed significant increases in melting temperature at the highest concentration of ligand (10 mM; 0.5°C for Asp and 0.3°C for Glu or 0.7°C and 0.6°C). Melting temperatures with ligand were compared with those in the absence of ligand by one-way ANOVA with Dunnett’s multiple comparison: ns p>0.05, * p<0.05, ** p<0.01, *** p<0.001, **** p<0.0001.

**Figure 9.**
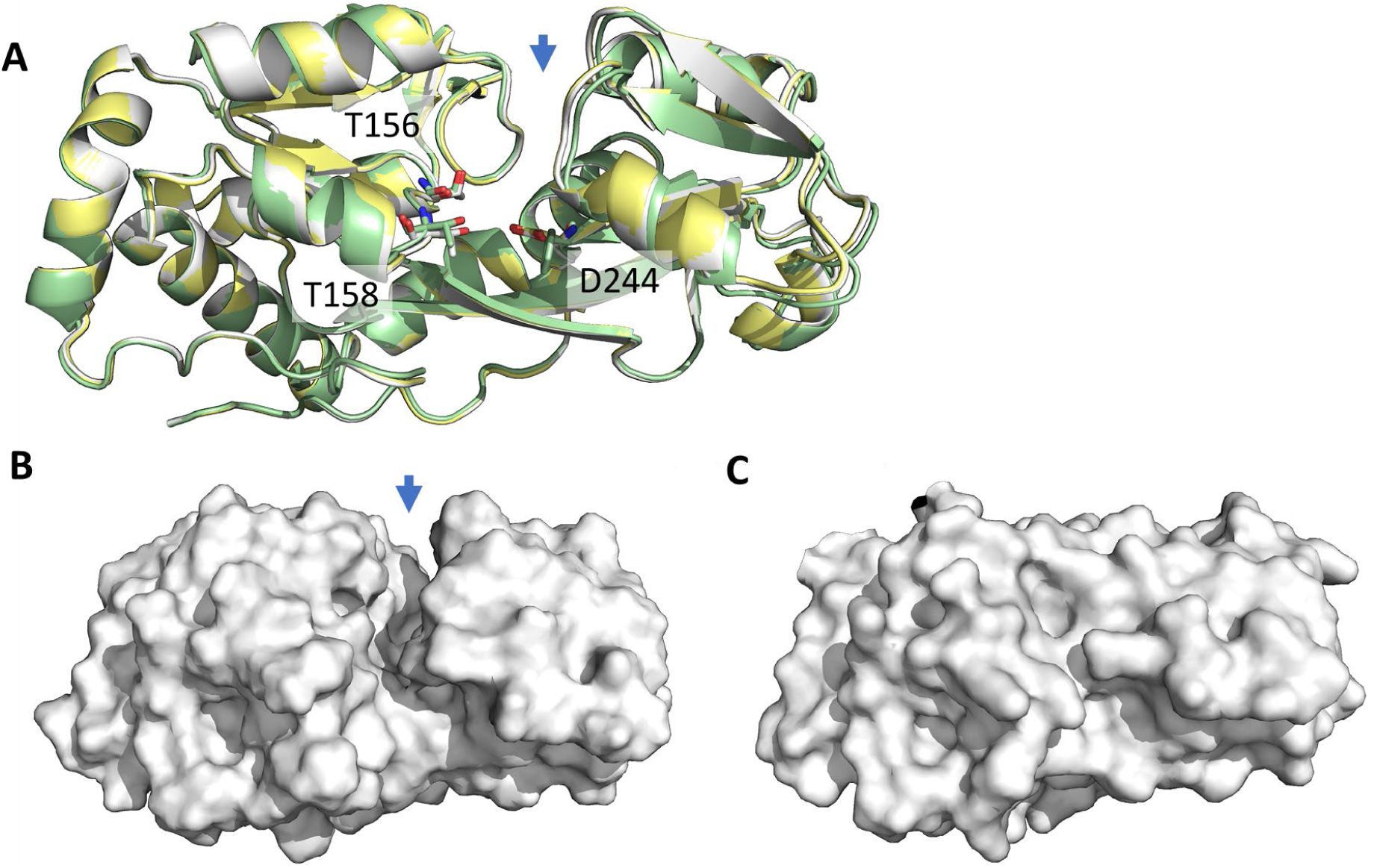
Apo GlnH had an open conformation with a pronounced cleft between the lobes. (A) Aligned crystal structures of Apo-GlnH_Ms_ (white) and variants GlnH_TA_ (yellow) and GlnH_DF_ (green) were determined at 1.58 Å, 1.64 Å and 1.60 Å. The blue arrow highlights a cleft between the two lobes, with solvent occupying the binding site. The structures of the mutants were similar to the wild type (R.M.S.D. values 0.098 and 0.458 Å respectively). Mutated residues are shown as sticks. (B) Surface views of Apo-GlnH_Ms_ and (C) Asp-bound GlnH_Ms_ show the presence of the cleft in the former but not the latter.

The lobes of the Apo structure were approximately 10 Å further apart, creating a cleft allowing access to the ligand binding site (fig 10 A & B). With the exception of Arg141, the side chains that contact aspartate retained their positions, forming hydrogen bonds with water (figure 10B). Arg141 rotated away from the ligand binding site and formed hydrogen bonds to to the side chain of Ser138 and two water molecules.

**Figure 10.**
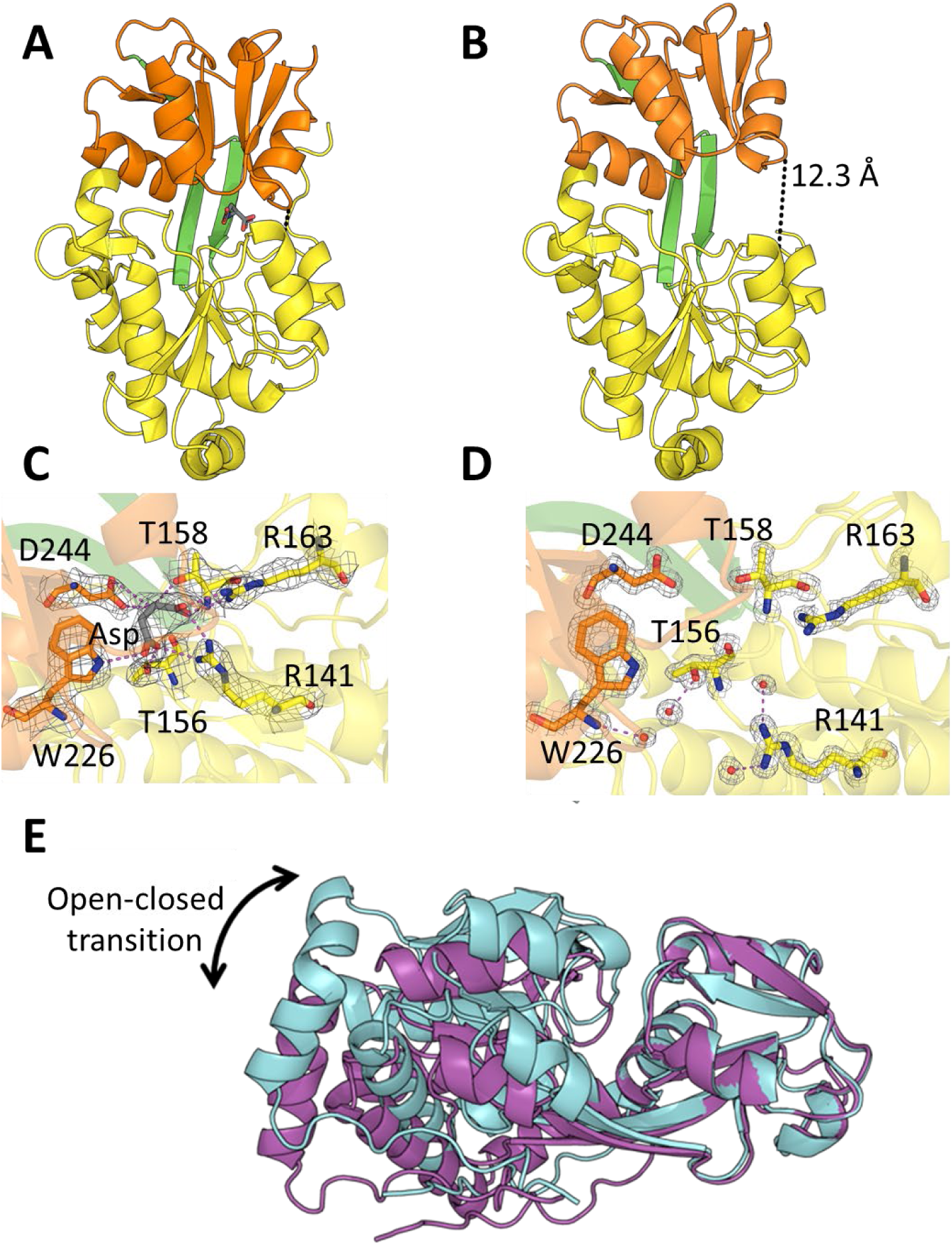
Comparison of aspartate-liganded and Apo GlnH_Ms_. (A&B) The large (yellow) and small (orange) lobes were approximately 8 Å further apart in the Apo conformation (A) than in the aspartate-bound conformation (B). (C&D) There was little rearrangement of the residues in the ligand binding site between the aspartate-bound (C) and Apo structure (D), with the exception of Arg141, which rotated away from the empty site. Waters in the cleft that hydrogen bond with these residues are shown as red spheres. (E) The aspartate-bound (blue) and Apo (purple) structures were aligned via the small lobe to highlight the protein movements involved in amino acid binding. The large lobe is rotated 10° in the open structure relative to the closed.

#### Effect of GlnH conformational changes on GlnX and PknG activity

To learn how the conformational changes in GlnH_Ms_ could impact its binding partner GlnX, AlphaFold 3 (13) was used to predict both the structure of GlnX and the interface between the two proteins (figure 11 A & B). The run produced five models, all sharing the same overall architecture and with high to moderate model confidence in all regions except for the N-terminus of GlnH, which would be membrane-anchored in the mature protein, and the N- and C- termini of GlnX (supplementary figure S7). The membrane topology of GlnX has been determined experimentally: the first 35 residues are cytoplasmic and followed by a transmembrane helix, a first periplasmic domain, a pair of transmembrane helices, and a second periplasmic domain and a final transmembrane helix (9). The periplasmic domains are predicted to form a pair of four-helix bundles that are aligned with each other and in contact such that the top (membrane distal) part of GlnX consists of eight alpha helices and with the four central helices contiguous with the transmembrane helices to form a club-like structure (figure 11 A). GlnH is predicted to bind at the top of the club, with the small lobe of GlnH predominantly contacting the first domain of GlnX, and the large lobe of GlnH predominantly contacting the second domain. Interestingly, the modelled conformation of GlnH is partway between the open and closed crystal structures, but closer to the closed liganded form. Because the modelled interface involves both lobes of GlnH, any change in the relative position of the lobes (due to ligand binding and release) would affect both domains of GlnX, consistent with its predicted role in signal transduction.

**Figure 11.**
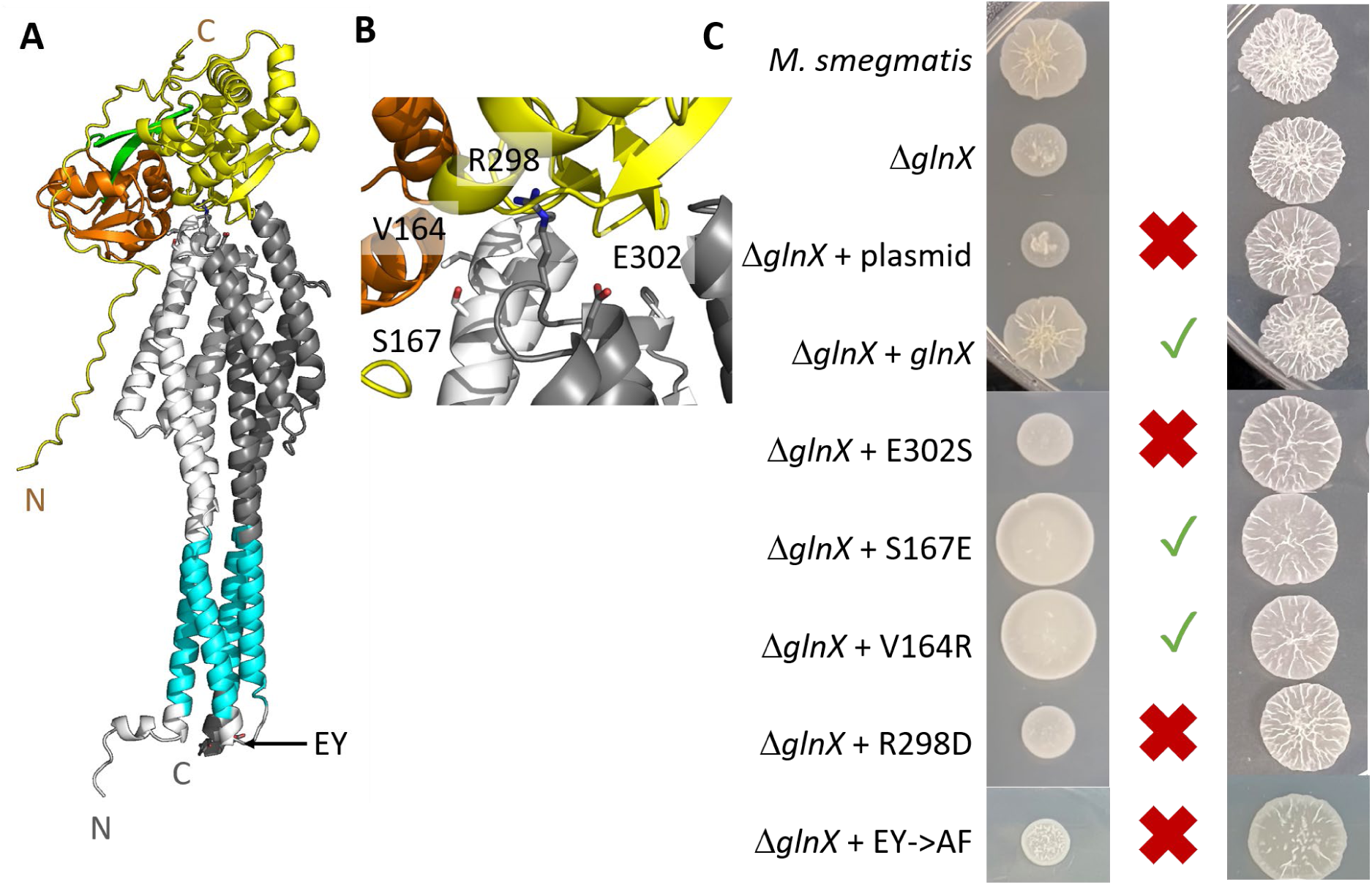
Investigation of the predicted GlnH-GlnX interface. (A) AlphaFold 3 was used to predict the structure of GlnX and model its interaction with GlnH_Ms_. The large and small lobes of GlnH_Ms_ are yellow and orange and the predicted lipoylated Cys is the first residue. GlnX is predicted to form two 4-helical periplasmic domains coloured light (domain 1) and dark grey (domain 2), connected by transmembrane helices. The first 16 residues of GlnX have been omitted as they were unstructured with a low confidence score. (B) V164, S167, R298 and E302 were predicted to lie within the interface of the AlphaFold model, and were selected for mutagenesis to characterise the interface. (C) GlnX residues highlighted in (B) were mutated and the mutants introduced into *glnX*-knockout *M. smegmatis* to determine whether they rescued the defect in glutamate utilisation. *glnX* knockout showed poorer growth than parent *M. smegmatis* on glutamate agar (left). Variants were scored by whether they restored normal growth (green tick) or failed to do so (red cross). All strains grew on standard mixed medium (Middlebrook 7H10 agar, right).

The model was analysed using PISA (14) to design mutations in the interface and test the requirement of the GlnH-GlnX interaction for regulating metabolism. A *glnX* knockout, already characterised in our laboratory, has a specific defect in glutamate utilisation, which mimics the *pknG* knockout (1). It is characterised by impaired growth on minimal media when using glutamate as the sole nitrogen source, but normal growth on standard rich media. Four separate mutations were introduced into the periplasmic part of GlnX. V164R and S167E are located in domain 1 and predicted to interact with the small lobe of GlnH, and E302S and R298D are in domain 2 and contact the large lobe of GlnH (figure 11 B, supplementary figure 8). Whilst wild-type GlnX complemented the *glnX* knockout, restoring growth on glutamate agar, variants E302S and R298D failed to complement this phenotype (figure 11 C & supplementary figures 9-10) despite normal levels of expression (supplementary figure S11), supporting the importance of the modelled interaction. The two other designed variants V164R and S167E were still able to complement the growth phenotype. To test putative interactions of GlnX with PknG, the C-terminus of GlnX was chosen for its high conservation (supplementary figure S8). A double mutant in the C-terminus E437A Y438F was constructed and failed to complement the *glnX* knockout (figure 11 C), despite normal expression (supplementary figure S9).

*pknG* and *glnX* deletion mutants have previously been described in *M. smegmatis* but *glnH* has not. To better understand the role of amino acid-binding in GlnH function, we constructed an in-frame deletion of *glnH*. Like the *glnX* knockout, the new *glnH* knockout grew normally in standard mixed medium (figure 12 A) but had slow, clumpy growth when glutamate was the sole nitrogen source (figure 12 B and C). The phenotype was restored by reintroduction of *glnH*. Variant GlnH_TA_ with low affinity for amino acids gave only partial complementation at 10 mM glutamate and no complementation at 1 mM glutamate (figure 12 E and F), consistent with GlnH acting as an amino acid sensor in the PknG pathway.

**Figure 12.**
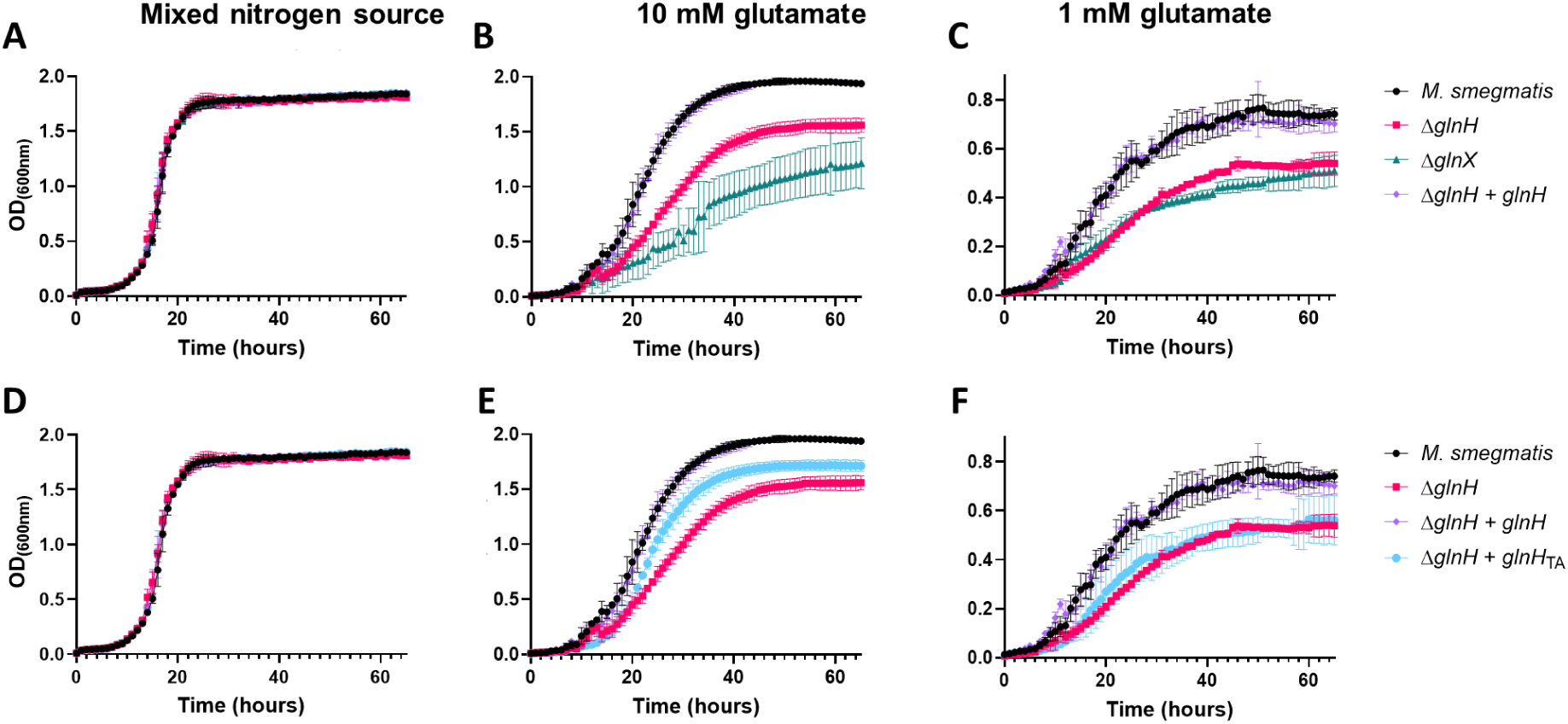
Deletion of *glnH* in *M. smegmatis* caused a growth defect that was complemented by GlnH_Ms_ but not by low affinity GlnH_TA_. Growth curves of *M. smegmatis* strains were performed in (A&D) standard mixed medium (Middlebrook 7H9), (B&E) minimal Roisin’s medium with 10 mM glutamate as the sole nitrogen source, and (C&F) minimal Roisin’s with 1 mM glutamate as the sole N-source. Graphs show the mean and standard deviation of 5 wells, and are representative of 3 independent experiments.

## Discussion

The new structure of the “open” Apo GlnH_Ms_ sheds the first light on how conformational changes of GlnH allow amino acid sensing. The opening and closing of GlnH in response to aspartate mirrors that of human glutamate receptors with a similar “Venus flytrap” fold, although the details of the conformational changes are different (15). The modelled GlnH-GlnX interface provides a rationale for how these conformational changes could be transduced by GlnX to modulate the activity of PknG. In particular, each lobe of GlnH is predicted to interact with a separate domain of GlnX, so the change in the relative position of GlnH upon ligand binding could impart changes to the relative positions of the domains of GlnX, transducing the signal across the membrane. Such a mechanism has been found for the bacterial amine sensor TmaT with its histidine kinase partner TmaS (16). However, different mechanisms have been found or proposed for other substrate binding protein sensors coupled to histidine kinases: the ligand may be required to initiate the interaction between sensor and transducer, or may cause the complex to dissociate (17). So far, the PknG system is the only characterised example of a SBP-stimulated serine threonine protein kinase. and further study will be required to discriminate between the possible mechanisms.

The defect in glutamate utilisation by *glnH* knockout *M. smegmatis* mirrored that previously reported for *glnX* and *pknG* knockout, consistent with the proposed signal pathway that regulates glutamate catabolism {Bhattacharyya, 2018 #1}, and has subsequently been linked to aggregation and biofilm formation {Varner, 2025 #49}. The failture of low-affinity GlnH to complement this phenotype provides the first direct evidence that amino acid-binding is a requirement for the function of GlnH. Interestingly, the growth defect of the *glnH* knockout was less severe than the *glnX* knockout (figure 11B). This difference probably reflects its upstream position within the PknG pathway. Thus, GlnX could potentially localise PknG to the periplasmic membrane for basal PknG activity, even in the absence of GlnH. By contrast, loss of GlnX removes the membrane tether of PknG, which lacks transmembrane domains, and so would reduce the likelihood of spontaneous PknG autophosphorylation in trans.

Many features of GlnH are conserved amongst the homologues: its overall structure and specificity for aspartate, and the ability for binding of aspartate to trigger conformational change of the large and small lobes with respect to each other. However, some details differed between *Corynebacterium* and *Mycobacterium*, or between pathogens and non-pathogens, most notably the affinity for aspartate, the conformational changes induced upon ligand binding, the oligomeric state, and the pH-profile of stability.

The affinity of GlnH for aspartate differed between *Mycobacterium* (K_D_ 0.23-0.57 μM) and *Corynebacterium* (264 μM). If the properties of recombinant GlnH reflect native membrane localised (lipoylated) GlnH then Mycobacteria would respond to nanomolar/low-micromolar concentrations of amino acids whereas *C. glutamicum* would only respond to high-micromolar/millimolar concentrations. Considering niches relevant to tuberculosis infection, extracellular concentrations of Asp and Glu are micromolar whereas cytoplasmic concentrations may be millimolar and phagosomal concentrations are presumed to be low due to nutrient limitation as an immune strategy (18–20). Although aspartate is the primary ligand for GlnH in terms of affinity (figure 1), glutamate is one of the most abundant free amino acids in many environments (20, 21) so may also be important in regulating PknG activity.

Structures of GlnH_Ms_ and GlnH_Cg_ were both determined in aspartate bound and free forms. However, the free-form of GlnH_Cg_ was of low resolution, so some caution must be taken in its interpretation. Nevertheless, the conformational changes in the two systems were different: in GlnH_Ms_ the lobes move apart in a hinge-like motion creating a cleft to allow access of ligand, whereas in GlnH_Cg_ the lobes rotate relative to one another about the long axis. Such a motion could still initiate signalling by imparting changes to the relative positions of the domains in GlnX and the molecular details of this process await further analysis.

Another difference between GlnH from *Mycobacterium* vs *Corynebacterium* is ligand-induced dimerisation of the latter but not the former. Bacterial substrate binding proteins are generally monomeric (7) and the dimerisation interface is only conserved in the closest homologues of GlnH_Cg_ (supplementary S6). Dimerisation is important in many signalling mechanisms, including serine threonine protein kinases of *M. tuberculosis* (22), but it is currently unknown whether dimerisation is involved in the PknG pathway. The GlnH_Cg_ homodimer is parallel with the lipoylated N-termini on the same face, so in principle is compatible with membrane anchoring. Interestingly, the predicted binding sites for GlnX are fully exposed in the homodimer. However, a membrane-anchored GlnH homodimer could not interact with GlnX in the bacterial membrane because the outermost promoter would be too far from the membrane surface (supplemental S12). Instead, GlnH dimers would have to first dissociate before binding to GlnX. Competition between dimerization and GlnX binding could potentially serve as an additional regulatory mechanism in *C. glutamicum* and its close homologues. It remains to be determined whether native, membrane anchored GlnH_Cg_ are able to dimerise.

GlnH differed between pathogens and non-pathogens in terms of its pH-dependence: GlnH from pathogens was stable across a broad pH range including pH 4-5, implying that it could sense amino acids in hostile and acidic microenvironments such as the phagosome or lung tissue and granulomas that may acidify due to enhanced glycolysis (23). Notably, the downstream effect of PknG activity is catabolism of glutamate (2). Catabolism of amino acids, particularly glutamate, is a key response for acid survival of several bacterial pathogens as it can raise pH locally by release of ammonia or bypass damaging or inhibited metabolic pathways (24–27). By contrast, GlnH from non-pathogens was more stable at neutral pH but unstable at low pH, presumably reflecting the native environments of the bacteria.

Conformational changes are an important mechanism of ligand sensing, but had not previously been characterised for this conserved and important Actinobacterial sensor. These results suggest amino acid-dependent conformational change of GlnH as a likely mechanism for amino acid sensing to modulate PknG activity and will require future studies of the transducer GlnX.

## Methods

### Production of recombinant GlnH

GlnH_Mm_ was amplified from genomic DNA of *M. marinum* M and inserted into an expression vector derived from pET43.1a(+) by ligation-independent cloning (primers in supplementary). GlnH_Ms_ was cloned from *M. smegmatis* mc^2^155 genomic DNA in a similar way, and GlnH_Mt_ and GlnH_Cg_ have previously been described (1). Proteins were expressed in *Escherichia coli* Shuffle® T7 express (New England Biolabs) grown in LB with 100 μg/ml ampicillin at 30°C with shaking and induced with 0.2 mM IPTG for 18 hours at 16°C. GlnH was purified by Ni-NTA affinity chromatography in Tris-buffered saline, followed by size exclusion chromatography in 10 mM Tris-HCl pH7.4, 150 mM NaCl then concentrated to 5-10 mg ml^-1^ using membrane concentrators (Vivaspin® Sartorius).

### Amino acid binding specificity and stability of GlnH

Amino acid binding and stability were determined using Thermal Melt Shift Assay. 25 μl reactions contained 0.15 mg/ml GlnH in 50 mM citrate-phosphate buffer pH 7 with 150 mM NaCl and a 1:10,000 dilution of SyPro Orange dye (Sigma-Aldrich). Amino acids were added to 10 mM unless otherwise indicated. The mixture was heated from 20-95 °C at a rate of 0.1 °C s^-1^ with measurement of fluorescence every 0.1 °C increment (QuantSutdio 5 qPCR system, Thermo-Fisher Scientific; TAMRA^TM^ reporter settings: excitation 552 nm, emission 578 nm). The melting temperature was the peak of the first differential of the curve of fluorescence vs temperature. Data plotted are the mean and standard deviation of triplicate measurements and statistical analysis used Graphpad Prism

### Isothermal calorimetry (ITC)

GlnH was dialysed three times for three hours against 1000-fold excess of phosphate buffered saline. Protein and aspartate solutions were prepared in dialysis buffer. ITC measurements were performed with a Malvern MicroCal PEAQ-ITC instrument at 25°C (GlnH_Ms_) or 35°C (GlnH_Mm_) using 300 μl of protein at 40 μM in the sample cell and aspartate (200 μM) in the syringe. Each run was started with an initial injection of 0. 4 μl followed by 12 injections of 3 μl each. Each experiment was performed in triplicate and data were analysed using Malvern MicroCal ITC analysis software with 1:1 binding stoichiometry to determine K_D_. Data are presented as mean ± 95% confidence interval.

### Site directed mutagenesis

*DpnI* mediated site-directed mutagenesis was used to create mutants of GlnH and GlnX using primer pairs listed in supplementary.

### Limited proteolysis using trypsin

Dialysed GlnH at a concentration of 0.3 mg/ml in 10 mM Tris-HCl pH 7.5 and 150 mM NaCl was incubated at for 1 hour with 0.02 mg/ml trypsin at 37 °C with addition of 10 mM Asp or dilutions of Asp (six 5-fold dilutios). Each reaction was stopped by boiling in 4X SDS loading buffer. The completeness of digestion was evaluated by SDS-PAGE (15%).

### Measuring oligomerisation of GlnH by size exclusion chromatography and mass photometry

Size exclusion chromatography used a superdex 200 10/300 column equilibrated with phosphate buffered saline with/without 10 mM Asp. Each run used 0.5 ml of dialysed GlnH at 40 μM and UV was monitored at 280 nm. Size standard proteins were run in parallel.

Mass photometry data was collected using a Two^MP^ instrument (Refeyn). Dialysed GlnH was diluted in PBS to 20 nM with or without 10 mM Asp. Data acquisition was performed using AcquireMP version 2024.1.1 (Refeyn) and was analysed using DiscoverMP version 2024.1.0 (Refeyn). A Gaussian curve was fitted to a contrast histogram (consisting of the number of detected molecules and their corresponding contract value) where each peak represents a subpopulation of molecules with a particular molecular mass. The contrast values of observed counts were converted into molecular mass by calibration using proteins of known molecular mass and size. The contrast-to-mass calibration curve was generated using NativeMark^TM^ Unstained Protein Standards (Invitrogen, Thermo Fisher Scientific). Default data acquisition settings were used according to the manufacturer (measurement mode, normal; image size, regular). Measurements were recorded for 60 seconds. Each sample was measured three times independently.

### Cryo-Electron Microscopy (Cryo-EM) grid preparation and data acquisition

3 µl of 0.68 mg/ml of Apo-GlnH in PBS, or 1.05 mg/ml GlnH in PBS plus 10 mM Asp, was applied to a glow-discharged (35 mA, 180 s; Quorum GloQube) holey gold grid (UltrAUFoil 1.2/1.3 300 mesh), followed by a 3 second blot plunge-frozen into liquid ethane using a Vitrobot mark IV (ThermoFisher Scientific) with the chamber kept at 4 °C and 100 % humidity.

All cryo-EM data were collected on a 300 kV Titan Krios ElectronMicroscope (Thermo Fisher) equipped with a Gatan K3 detector, with EPU data acquisition software (ThermoFisher Scientific) at the Midlands Regional Cryo-EM Facility, University of Leicester. Data collection parameters are provided in supplementary table 2.

### Cryo-EM image processing

All single particle cryo-EM datasets were processed in cryoSPARC (28). In short, CTF correction, motion correction, and particle blob picking were first performed. Particles were extracted using a box size of 256 and 180 pixels for the Asp-bound homodimer and Apo-monomer, respectively. Particles were selected and optimised through iterative rounds of 2D classification, ab-initio reconstruction, heterogeneous refinement and 3D classification without alignment. The best maps were then further refined using homogenous refinement and finally non-uniform refinement. The cryo-EM processing workflow are shown in supplementary figure 3 for aspartate-bound homodimeric GlnH_Cg_ and supplementary figure 4 for apo-GlnH_Cg._

### Cryo-EM structure refinement and model building

Post-processing sharpening on cryo-EM maps was performed prior to modelling. Maps were sharpened in Phenix with Auto_sharpen (29) and using DeepEMhancer (30). A model of GlnH_Cg_ was generated using AlphaFold3 (13) and rigid-body fitted into the cryo-EM density maps in UCSF chimera (31). Iterative manual model building and refinement was performed in COOT (Emsley *et al.,* 2010) and PHENIX (32). Refinement statistics are provided in supplementary table 2.

### Crystallisation of GlnH

Crystallisation trials were performed using commercial screens using the sitting drop vapour diffusion method in 96-well plates at 22°C. GlnH_Mm_ crystallised in 0.1 M Tris-HCl pH 8, containing 0.04 M manganese chloride and 18% (w/v) PEG 8000. GlnH_Ms_ crystallised in 0.1 M Tris-HCl pH 8.5, 0.2 M sodium acetate, 0.01 M sodium aspartate, 4% (v/v) 1,3-propanediol, 30% (w/v) PEG 3350. GlnH_TA_, GlnHDF and unliganded GlnH_Ms_ crystallised in 0.1 M sodium acetate pH 5 containing 0.2 M ammonium chloride and 20% (w/v) PEG 6000.

Diffraction data were collected on the beamline at Diamond Light Source and processed with Xia2.DIALS (. All diffraction datasets were processed within the CCP4i suite and PHENIX (32–34). Phases were determined by molecular replacement with Phaser, using the Asp-liganded structure of GlnH_Mt_ (6H1U) as a search model. Models were optimised using iterative manual model building and refinement in COOT and PHENIX. Data collection and refinement are provided in supplementary table 3.

### Functional characterisation of GlnX variants

*glnX*-deleted *M. smegmatis* mc^2^ 155 (Δ*glnX*) was previously described (1) and was routinely cultured at 37 C using Middlebrook 7H9 broth with 10% ADN (0.5% bovine serum albumin, 0.2% dextrose, 0.085% NaCl) and 0.05% v/v tween 80, or on Middlebrook 7H10 agar with 10% ADN. When needed, hygromycin was included at 200 μg/ml. To assess utilisation of glutamate as the sole nitrogen source, cultures were grown to mid-exponential phase and spotted onto Sauton’s agar 3.7 mM KH_2_PO_4_, 2 mM MgSO_4_, 9.5 mM sodium citrate, 0.17 μM ferric ammonium citrate, pH 7.0, 1% glycerol and 10 mM glutamate.

A new complementation plasmid was constructed to include a Strep tag to allow detection of expression level by Western blot. *glnX* was amplified by PCR from *M. smegmatis* genomic DNA and cloned using In-Fusion kit (Takara Bio) in shuttle plasmid pMyNT for Mycobacterial expression (hygromycin selection, acetamide induction)(35). Plasmids were introduced to *M. smegmatis* Δ*glnX* by electroporation.

To detect expression of recombinant *glnX*, the relevant strain of *M. smegmatis* was cultured in 10 ml Middlebrook broth to exponential phase, washed with PBS then lysed by bead-beating in PBS. Lysate was analysed by SDS PAGE (to normalise protein yield) and Western blot using Strep•Tag® II Antibody HRP Conjugate (Novagen) followed by SignalFire^TM^ ECL Detection Reagent (Cell Signalling Technology).

### Deletion of *glnH* from *M. smegmatis* to generate *ΔglnH*

An in-frame, unmarked *glnH* deletion mutant was constructed by homologous recombination using a published method (36). Briefly, a suicide plasmid was assembled using InFusion (Takara BioSciences) from three PCR amplicons: the upstream and downstream regions from *glnH* amplified from *M. smegmatis* genomic DNA and p2NIL. The marker gene cassette was added as described (36). Initial transformants were selected on kanamycin (30 μg/ml). Counterselection used 2% sucrose and putative double crossover unmarked deletion mutants were verified by PCR. A complementation vector was constructed by cloning *M. smegmatis glnH* into pMV306 with the cognate *glnX* promoter (1).

Growth curves of *M. smegmatis* were carried out in 96-well plates at 37C with shaking, in Middlebrook 7H9 broth (control) or minimal Roisin’s containing 10 mM or 1 mM glutamate as the sole nitrogen source, and 0.4% glycerol plus 0.05% Tween 80 as carbon sources.

## Acknowledgements

We acknowledge The Midlands Regional cryo EM Facility at the Leicester Institute of Structural and Chemical Biology (LISCB), major funding from MRC (MC_PC_17136). We thank Diamond Light Source (DLS) for access to beamline I04 via the UK Midlands Block Allocation Group mx34438. We also thank beamline scientists at DLS for help with data collection. pMyNT was a gift from Annabel Parret & Matthias Wilmanns (Addgene plasmid # 42191; http://n2t.net/addgene:42191; RRID:Addgene_42191). We would like to thank J Basran for assistance with ITC.

This work was supported by the University of Leicester Future 100 Scholarship (HLT) and the MRC grant MC_PC_17136 (AKC). The funders had no role in study design, data collection and analysis, decision to publish, or preparation of the manuscript.

